# Cooperativity boosts affinity and specificity of proteins with multiple RNA-binding domains

**DOI:** 10.1101/2021.01.27.428308

**Authors:** Simon H. Stitzinger, Salma Sohrabi-Jahromi, Johannes Söding

## Abstract

Numerous cellular processes rely on the binding of proteins with high affinity to specific sets of RNAs. Yet most RNA binding domains display low specificity and affinity, to the extent that for most RNA-binding domains, the enrichment of the best binding motif measured by high-throughput RNA SELEX or RNA bind-n-seq is usually below 10-fold, dramatically lower than that of DNA-binding domains. Here, we develop a thermodynamic model to predict the binding affinity for proteins with any number of RNA-binding domains given the affinities of their isolated domains. For the four proteins in which affinities for individual domains have been measured the model predictions are in good agreement with experimental values. The model gives insight into how proteins with multiple RNA-binding domains can reach affinities and specificities orders of magnitude higher than their individual domains. Our results contribute towards resolving the conundrum of missing specificity and affinity of RNA binding proteins and underscore the need for bioinformatic methods that can learn models for multi-domain RNA binding proteins from high-throughput *in-vitro* and *in-vivo* experiments.

## Introduction

RNA-binding proteins (RBPs) regulate various steps of mRNA biogenesis including RNA splicing, localization, translation, and degradation [1]. In order to ensure that these proteins bind the correct set of RNA molecules, the interactions have to be highly specific. This can be achieved through cooperative binding, in which two or more relatively weakly binding RNA-binding domains (RBDs) are combined, to form one high affinity complex [2, 3]. Around 45% of eukaryotic RBPs have at least two binding domains [4] (Figure 1). In addition, these domains often recognize very short and degenerate RNA motifs (~ 3, rarely more than 5 nucleotides) [5, 6], while typical DNA-binding domains bind sequences of around 6 to 12 nucleotides [7, 8, 9]. Because of their larger binding sites, the dissociation constants (*K*_d_) of most transcription factors are in the picomolar to nanomolar range, while individual RBDs often only bind with affinities in the micromolar to millimolar range.

**Figure 1:**
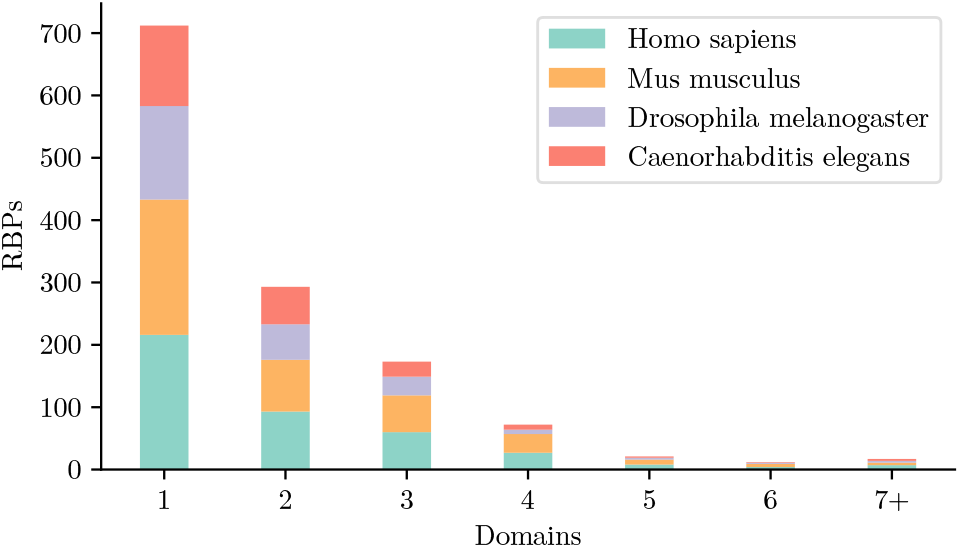
Many RBPs have more than one domain. Numbers of RNA binding domains per protein. The data in the graph was taken from the RNA-binding protein database (RBPDB) [4] and shows how many RBDs every protein in this database has. It includes proteins from human, mouse, Drosophila and Caenorhabditis elegans.

RBPs use multi-domain RNA recognition to bind with high affinity and specificity, despite the low affinity of their individual domains. The affinity increase in cooperative binding stems from an increased local concentration of the RNA molecule at an unbound binding domain when another domain is already bound [10]. The neighboring RNA binding site is kept closer to the second RBD since the molecules are already linked. This adds to the background concentration, resulting in a much higher effective concentration *c*_eff_. This high local concentration in turn increases the rate for the binding reaction. The concept of an effective concentration has been introduced before [10, 11], and was used to thermodynamically model different systems with two binding sites, based on the interactions of the isolated sites [11, 12, 13, 14].

A general quantitative model of cooperative multi-domain interactions can be adapted to systems other than RNA binding. Cooperative binding is also utilized by antibodies [10, 13], DNA-binding proteins [12] and many components of phase-separated biological droplets [15, 16]. In all of these cases, the combination of multiple binding sites and their connection through flexible linkers increases binding affinity in a completely analogous way to multi-domain RNA binding.

For all these cooperative interactions it is important to understand the binding affinity of the complex. So far, existing models have only described cooperative binding between two domains, with flexible linkers between the domains of one binding partner. The concentration enhancing effects of flexible linkers were either modelled with a uniform distribution in a sphere with radius *L* around the first domain, where *L* is the length of the chain. Other approaches used the more detailed worm-like chain model.

Here, we develop an equilibrium thermodynamic model for cooperative RNA-protein interactions and use the worm-like chain model for the behavior of flexible polymer chains. We validate our analytical calculations with stochastic simulations using Gillespie’s algorithm [17, 18]. Using our model, we can show that the dissociation constant decreases exponentially with each added binding site, by which very high affinities and specificities can be achieved with low-affinity and -specificity RNA binding domains. Our calculations can be extended to any number of binding sites and are able to also include situations, in which both binding partners contain flexible linkers between binding sites. We compare predictions of our model to four proteins, for which the affinities of individual domains have been measured and show that our estimated *K*_d_s are in good agreement with the experimental values.

## Material and Methods

We will first describe our model and assumptions. We will then illustrate how the dissociation constant for a protein with two or three RNA binding domains can be derived, and generalize the derivation to any number of binding domains.

### Simple cooperative binding model

The model describes the cooperative, multivalent binding of RNA-binding proteins possessing *N* RNA-binding domains to an RNA with *N* binding sites (Figure 2A). To be able to analytically calculate the effective dissociation constant for the proteins and its RNA substrate, we need to make two simplifying assumptions. First, we assume that each RNA-binding domain can only bind to a single, cognate binding site on the RNA, so domain 1 to RNA site 1, domain 2 to RNA site 2, and so on. We further assume that an RNA is at most bound by a single protein. This is good approximation as long as the local concentration of domains of the already bound protein at the RNA sites is much larger than the background protein concentration. In that regime, the first-bound protein will outcompete all other proteins from binding to its protein.

**Figure 2:**
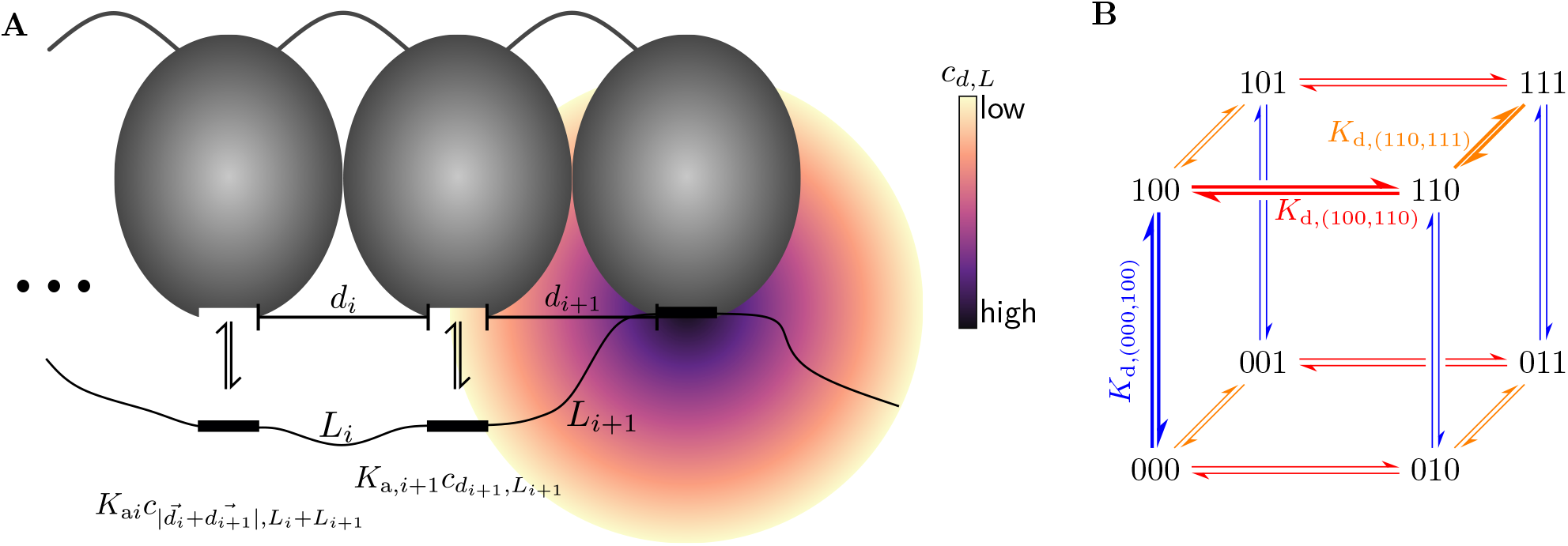
Thermodynamic model for cooperative RNA-protein interactions. **(A)** Illustration of an RNA with an RNA-binding protein (RBP) binding to it. All binding sites on the RNA are only bound by one domain of the RBD. Each of these interactions has its individual *K*_d_. The worm like chain model is used to estimate the concentration *c_d,L_* of an RNA-binding domain at an RNA binding site when another domain at distance *d* in the protein is already bound to the RNA a linker length *L* away. **(B)** Representation of the reaction network at the example of three binding sites on the RNA. Every possible reaction step has a dissociation constant equal to the individual *K*_d_ for the domain-to-RNA-site interaction times the concentration of the domain at its cognate site (eq. 8).

The transitions in this model from one binding configuration to the next can be described by a coupled reaction system with 2^*N*^ different binding configurations, indexed by a binary string that indicates which sites are bound, e.g. (101) represents the configuration in which the first and third sites on the RNA are bound. Figure 2B shows the reaction network for *N* = 3 illustrated as a cube. The dissociation constants of the individual binding steps are called *K*_d,(*x*,*y*)_, for a reaction between configuration *x* and configuration *y*. The effective concentration of the RNA-binding domain at another RNA site when one domain is already bound to its site is denoted as 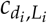, with *d_i_* the distance between domains and *L_i_* the chain length between RNA binding sites. For example if we consider the reaction 001 ⇌ 011 we would write the equilibrium constant as

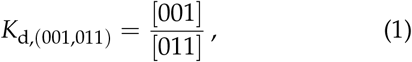

and the concentration at the second binding site as 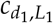.

### Effective concentrations using the worm-like chain model

To estimate the effective concentration of a protein domain at an RNA binding site if another domain of the protein already binds the RNA we model the RNA and the disordered peptide linker between RNA-binding domains as worm-like chains [19]. We use estimates of the persistence length *l*_p_ of RNA and disordered peptides, and we call *L* the chain length between RNA-binding sites. The probability density at an end-to-end distance of *d*, where *d* is the distance between binding domains on the protein, can be interpreted as the local concentration of the domain at its RNA binding site, as seen in figure 2A. Given a large enough chain length between the binding sites, *L* ≫ *l*_p_, the radial distribution function tends towards a Gaussian distribution,

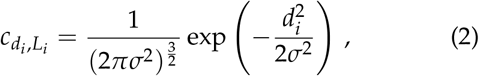

with a variance of

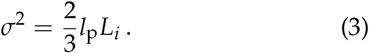

This assumes a rigid connection between protein domains. If the protein has flexible linkers between the binding domains, allowing them to move independently, we get (see Supplementary Methods for a more detailed derivation)

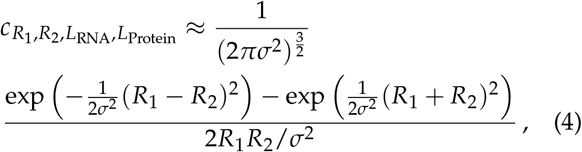

with a variance of

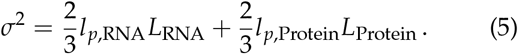

### Calculating K_d_ values for two or three domains using equilibrium thermodynamics

As shown before, the total dissociation constant for a protein with two domains binding to a chain with two binding sites can be written in terms of association constants of its individual domains as

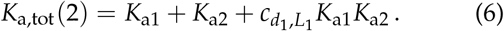

To get some intuition for this equation, we consider the law of mass action for any binding step, when at least one domain is already bound, for example 01 ⇌ 11:

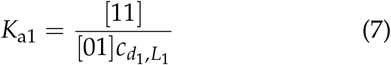

and by rearranging we get

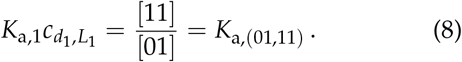

Thus, equation 6 can also be written as

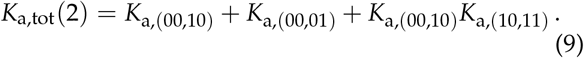

In the Supplementary Methods we derive the *K*_d_ for *N* = 2 for the alternative case where the two RNA-binding domains have the same specificities such that each of them can bind to any of the two binding motifs on the RNA.

To extend the previous treatment to three binding sites, we write the total dissociation constant *K*_d_ in terms of the individual concentrations of all binding configurations. Simple rearranging leads to the *K*_d_ in terms of dissociation constants of the individual domains (Supplementary Methods),

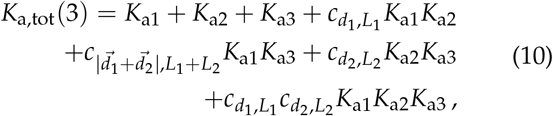

or in a more general notation

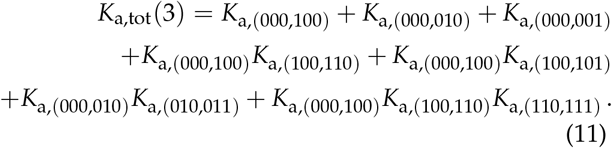

### Calculating K_d_ values for N binding sites

To generalize this approach to *N* binding sites, we can similarly express the observed dissociation constant according to the law of mass action in terms of the equilibrium concentrations of all states:

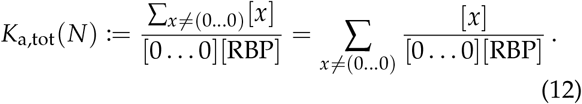

[*x*] denotes here the concentration of the *x*’th state, [0 … 0] is the concentration of the unbound RNA and [RBP] the concentration of the unbound protein. The individual terms can be interpreted as apparent association constants for a theoretical one-step reaction from the unbound state to state *x*, so that

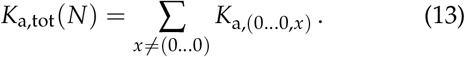

The *K*_a,(0…0,*x*)_ can be written as a product

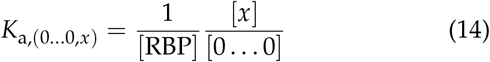

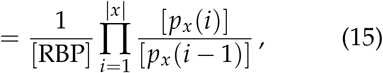

where |*x*| is the number of bound sites at state *x* and *p_x_* describes a path from the unbound state (0 … 0) to state *x* = *p_x_*(|*x*|). We define 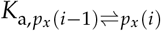 as the association constant of the reaction between configurations *p_x_*(*i* − 1) and *p_x_*(*i*) along the path *p_x_* (Figure 2B). Because for the first factor we can write

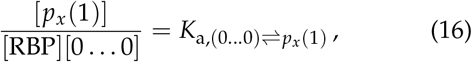

equation (15) can finally be written as

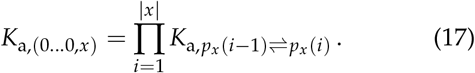

It can readily be seen that

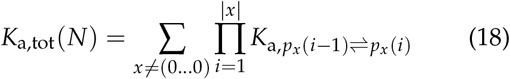

is a generalization of equations (10) and (6).

If the individual *K*_a_s are large, the last term of the sum dominates

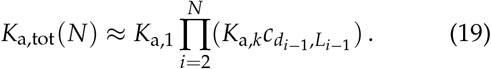

This shows how each added binding site approximately multiplies the total *K*_a_ by a factor 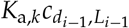.

### Simulation of cooperative binding with Gillespie algorithm

We validated our analytical calculations, described above using the Gillespie algorithm [17, 18] implemented in the Python library Gillespy2 [20]. We performed simulations of our model by defining all binding configurations as molecular entities in the simulation and determining the *K*_d_ based on trajectories of the systems (Supplementary methods).

### Determining the model parameters

*K*_d_ values of individual binding domains are taken from experimental measurements like electrophoretic mobility shift assays (EMSA) or isothermal titration calorimetries (ITC). Distances between binding sites on the protein are Euclidian distances calculated based on available PDB structures. The contour lengths of ssRNA linkers between binding sites and the length of flexible linkers between protein domains are estimated as the number of nucleotides or amino acids multiplied with a length per base of 5.5 Å (mean of 5 measurements) [21, 22, 23, 24, 25] or a length per aminoacid of 3.8 Å [26] respectively. The persistence length *l*_p_ of ssRNA and disordered proteins is estimated as 2.7 nm (mean of 5 measurements) [21, 22, 23, 24, 25] and 3.04 Å [26] respectively.

## Results

### The thermodynamic model correctly estimates measured dissociation constants

To validate our model, we estimated the dissociation constants for several RBPs by performing simulations and using our analytical approach (Figure 3). For these proteins the *K*_d_ values for individual domains and the whole protein have been measured experimentally. The proteins used were the zipcode binding protein 1 (ZBP1) [27], the heterogeneous nuclear ribonucleoprotein A1 (hnRNP A1) [28], the two terminal domains of the polypyrimidine tract binding protein (PTB) [29, 30] and the first four domains of the insulin-like growth factor 2 mRNA-binding protein 3 (IMP3 or IGF2BP3) [31]. The first three of these proteins consist of two rigidly linked domains. In contrast, IMP3 consists of three domain pairs with flexible linkers between the pairs. In our model the first two of the three IMP3 domain pairs were represented as two binding sites, connected by a flexible linker. All predictions were at least within a factor ~ 5 of the experimental value (see Supplementary Information), demonstrating the applicability of the model to multivalent, cooperative binding of RBDs to their RNA substrates.

**Figure 3:**
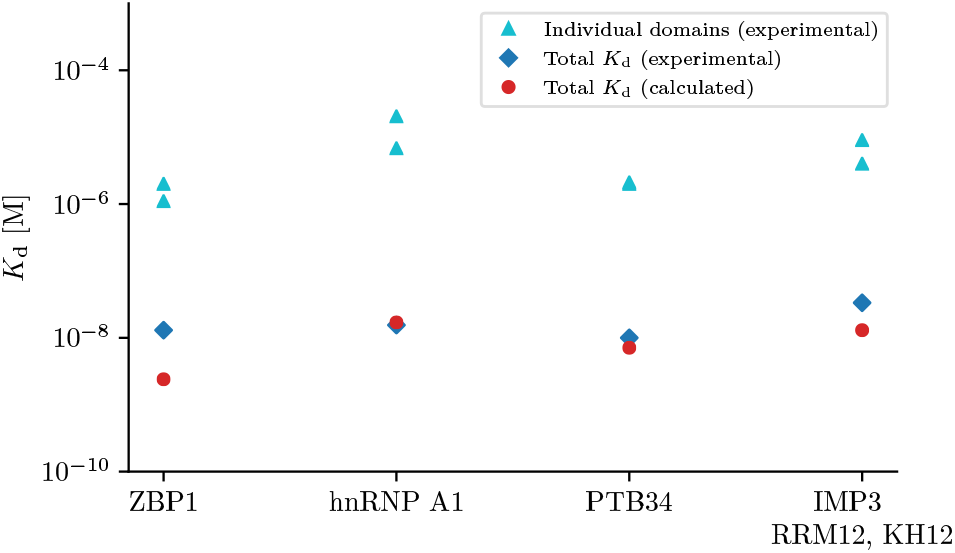
Measured *K*_d_s are in good agreement with model predictions. We found four RBPs composed of two RBDs for which dissociation constants of the full-length protein had been measured together with those of individual RBDs [27, 28, 29, 30, 31]. We used the simple thermodynamic model to estimate the effective *K*_d_’s of the full-length RBPs from those of their individual domains and from linker lengths and protein binding site distances *d* and found agreement within a factor of ~5. No free fitting parameters were used (See Supplementary Methods for details).

### K_d_ decreases exponentially with the number of binding sites

We then asked how the effective dissociation constants for RBPs depends on the number *N* of their RBDs (Figure 4A). We chose *K_d_* values for RBDs and linker lengths in the ranges of typical RBPs. We observed an exponential increase in affinity with the number of binding sites with an incresase by a factor *K*_a,*k*_ *c*_*d,L*_ for each added domain (equation 19). The local concentration of the RBDs, *c_d,L_*, depends on the linker length *L* between consecutive binding sites and the distance *d* between the consecutive RBDs as shown in equation (2). While the factor in real RBPs will depend on individual *K*_d_s and distances between binding sites, the analysis shows that the global *K*_d_ can drop by orders of magnitude per domain added.

**Figure 4:**
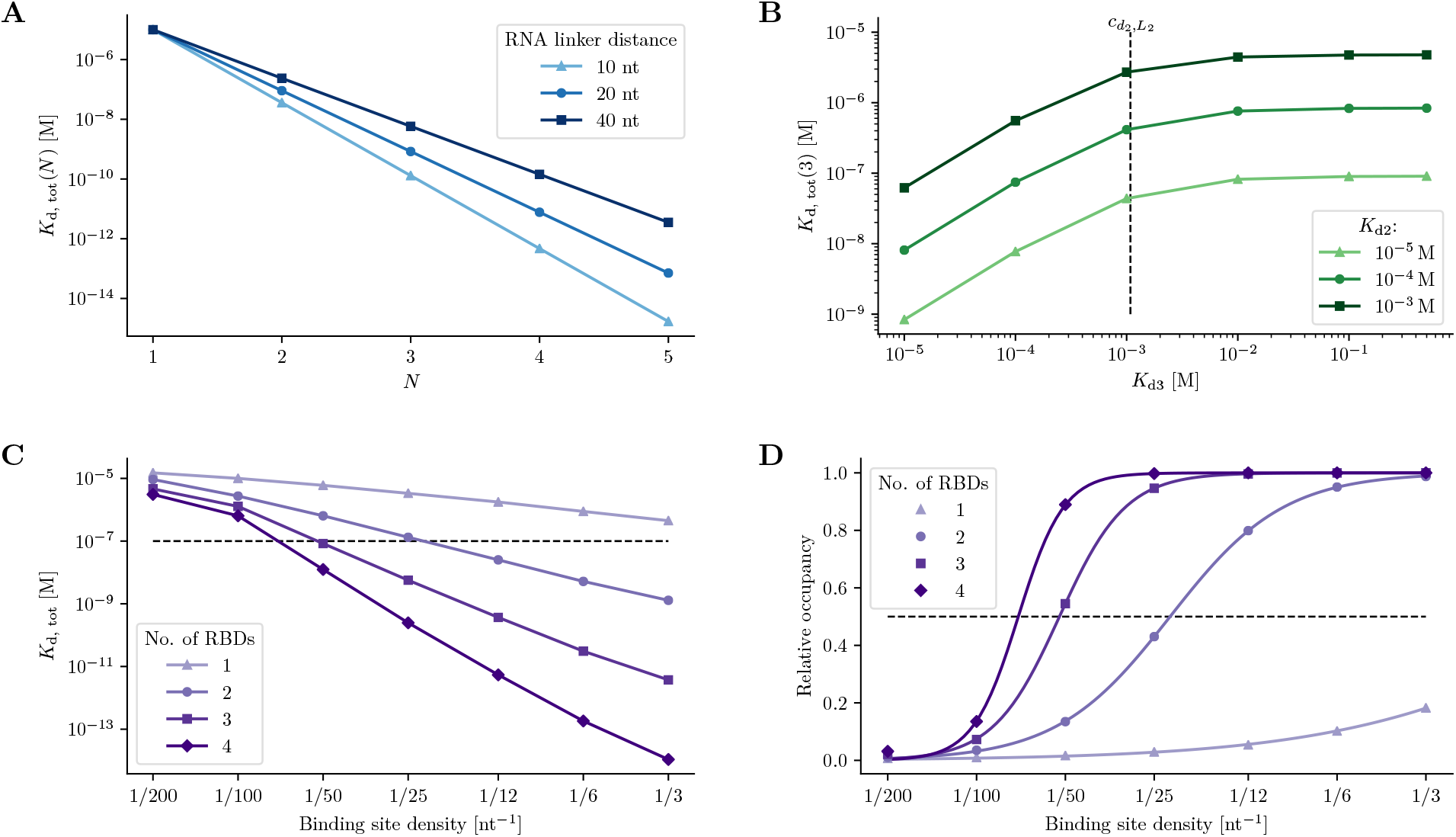
Dependence of the effective dissociation constants of RBPs on the number and *K*_d_’s of RBDs, the effective concentration *c_d,L_*, and the binding site density on the RNA. **(A)** Dissociation constants decrease exponentially with the number of binding domains. All RBD *K*_d_’s were set to 10 μM and distances *d* between rigidly linked binding domains to 2 nm. The relationship between the number of binding sites and the *K*_d, tot_ is shown for RNA binding sites spaced at 10nt , 20nt, or 40nt. **(B)** Individual RBDs contribute to the total *K*_d_ up to a threshold *c_d,L_* in the individual *K*_d_. In the calculations we used a *K*_d_ of 10 *μ*M for the first domain. The *K*_d_s for the second and third domain are *K*_d,2_ and *K*_d,3_, as indicated in the plot. This was calculated for equal distances between rigidly linked binding domains of 2 nm and an RNA linker length of 20 nt. **(C)** Dissociation constants decrease with the binding site density on the RNA. As a simplification we assumed that the domains bind sequentially the binding sites on the RNA. *K*_d_s of individual domains were 100 μM and the total RNA length was 200 nt. The horizontal line indicates a concentration of 0.1 μM used for the calculations in Figure D. **(D)** Relative occupancy of the RNA as measured by [RNA]/(*K*_d_ + [RNA]) as a function of binding site density on the RNA at an RNA concentration of 0.1 μM (horizontal line in Figure C). Curves show fits with sigmoidal Hill-functions, with Hill coefficients of *n*_1_ = 0.96, *n*_2_ = 2.26, *n*_3_ = 3.88 and *n*_4_ = 5.65 for one to four domains, respectively.

### Contributions of individual domains to the total K_d_ becomes negligible after a threshold in the individual K_d_

To further investigate the effect of domain *K*_d_s to the total affinity, we calculated the dissociation constants for artificial RBPs with 3 domains, kept the *K*_d_ of the first domain constant and varied *K*_d,2_ and *K*_d,3_ (Figure 4B). As expected, the total *K*_d_ increases when the *K*_d_ of one individual domain is increased. According to equation (18), when 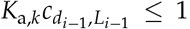, or, equivalently, 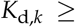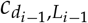, the contribution of domain *k* to the total *K*_d_ quickly saturates (horizontal line in Fig. 4B).

### Multidomain RNA binding proteins can achieve high specificity by modularity and a varying binding site density

It is known that the combination of multiple RNA binding domains is important for providing the specificity needed to bind to the correct target RNAs [32]. The density of binding sites on the RNA molecule is also an important indicator of binding affinity and specificity [33]. To investigate this effect, we calculated the *K*_d_ and the relative occupancy of RBPs as a function of the binding site density on the RNA (Figure 4C and D). To simplify, we assumed that the domains always bind sequential binding sites on the RNA. Intuitively, the occupancy increases with increasing binding site density for all numbers of protein domains. In proteins with multiple domains, however, the transition in the binding curve is much more rapid than for proteins with one domain, allowing high levels of occupancy even at low binding site densities. To quantify the cooperativity, we fitted a sigmoidal function 1/(1 + (*D*/*D*_0_)^*h*^) with binding site density on the RNA of *D* and Hill coefficient *h*. We observe that the Hill coefficient, which is a common measure of cooperativity, is close to the number of RBDs, although growing somewhat faster (*n*_1_ = 0.96, *n*_2_ = 2.26, *n*_3_ = 3.88 and *n*_4_ = 5.65 for one, two, three and four domains, respectively).

## Discussion

### Simplified assumptions can limit model accuracy

Our estimated *K*_d_ deviates to some extent from the experimental measurement (Figure 3). Some simplifications in the model can potentially explain these differences. Most notably a lot of linkers between RNA binding sites are very short. To estimate *c*_eff_ we use a Gaussian distribution, which is valid if the chain length is much larger than the persistence length of the RNA, *L* ≫ *l*_p_. If the chain length is shorter, the end-to-end distribution will not be radially symmetric anymore, but depend on the initial tangent orientation of the bound end [34]. It has been shown that only for 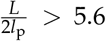 the distribution has a single maximum near the origin in direction of the initial orientation and approaches a Gaussian for larger values [34]. The chain lengths in the examples given earlier correspond to rather stiff chains. Depending on the orientation of the next binding site in relation to the first, the effective concentration and consequently the final *K*_d_ can be overestimated or underestimated. In order to increase the accuracy of estimates for *c*_eff_ one would have to take into account other geometric properties of the protein next to the distance between binding sites. However, for short-range systems, the analytical solution to the worm-like chain becomes highly complex and with alternatives like molecular dynamics simulations the wanted simplicity and intuition of the model is likely to be lost.

In addition to the short RNA linkers, RNA secondary structure and unspecific binding might also play a role in decreasing accuracy of the predictions. Furthermore, the sequence of the RNA also influences its flexibility. A lot of measurements of the persistence length of ssRNA are being done with very repetitive sequences. This effect will most likely average out for long chains, but can still be present in short ranges considered here.

### Thermodynamic model extends previous models of cooperative binding

The described model allows to estimate the thermodynamic behavior of cooperative interactions for any number of binding sites. There are existing quantitative descriptions of cooperative binding. Crothers and Metzger developed a model to determine the overall *K*_d_ of the two binding sites of an antibody, estimating *c*_eff_ with the particle-in-a-sphere model, assuming, that the RNA binding site is uniformly distributed inside a sphere with a radius of *L* around the first already bound binding site [10]. This model has been extended several times, taking into account different properties like chain length of the flexible linker between binding sites/domains and also transferring it into the context of RNA binding [11, 12, 13, 14]. All of these studies, however, derive dissociation constants for two domains. The results for *N* = 2 match our model and have now been extended to an arbitrary number of binding sites.

In addition, previous models have only described flexible linkers between binding sites on one binding partner. Since both the RNA is a semiflexible polymer and there are many examples of proteins with flexible linkers between the domains, this has been extended in our model to allow flexible linkers on both binding partners.

### Other mechanisms for increasing affinity and specificity

Our model presented here assumes that RBPs bind to well defined binding sites. In many cases, however, the short and degenerate motifs on the RNA can be repeated, creating the possibility for multiple binding registers [2, 3, 35]. This can be seen for example in the HuR C-terminal RRM binding to AU-rich RNA regions [36] and also in PTB, one of our examples, which binds to polypyrimidine tracts [37]. When the repeat regions are long enough, the proteins can bind in more than one arrangement. The effects on the affinity by encompassing *n* binding registers in the RNA motif can be estimated through a simple statistical consideration by a linear factor of *n*.

Moreover, the concept of “fuzziness” has been qualitatively described by Olsen et al. [38]. It describes the more general situation when every RNA binding site can at least to some degree bind to every protein domain. We calculate this effect in our model for two binding sites (Supplementary Methods). Including fuzzy binding in the calculations increases the number of possible bound configurations and thus the complexity of the combinatorics. However, it does not qualitatively change the results that we present here.

### Multidomain RNA binding can promote phase separation

It is well known that phase-separated biological droplets, which function to concentrate and organize molecules inside the cell, form multivalent networks. These multivalent interactions can arise from weak interactions between intrinsically disordered regions of the proteins and/or by multivalency through multiple connected domains [16]. We are, however, still lacking a good quantitative description of the underlying mechanisms behind phase separation in biology. Many stages of RNA metabolism also involve phase separation [16, 39, 40]. These condensates that form around RNA molecules have been called scaffolded condensates [41, 42]. Understanding cooperative multidomain binding in the simple case described here, could also give some insight into the formation of phase-separated RNA-protein complexes.

## Conclusion

We have shown here a general method for evaluating the affinity of multivalent cooperative binding interactions based on the affinity of the individual non-cooperative interactions. This model underlines the large importance of cooperativity in multivalent binding. It demonstrates how highly specific and affine interactions can be made possible even at very low concentrations. Besides RNA-protein binding this model can be applied to many different systems such as antibodies or the binding of proteins to intrinsically disordered regions in proteins.

## Code and data availability

The code used for the simulations and calculations is available at https://github.com/soedinglab/cooperative_rbp. The protein structures used in the validation of the model are available under the PDB accession codes 2n8l, 6dcl, 2adc, 6fq1 and 6gqe.

## Contributions

SHS implemented the algorithms and conducted all the analysis. JS conceptualized the idea. SSJ and JS supervised research. SHS, SSJ, and JS wrote the manuscript.

