## Supplementary Material for "Cooperativity boosts affinity and specificity of proteins with multiple RNA-binding domains"


#### 1 Procedure for calculating a dissociation constant for $N = 3$

We describe an approach to calculate the dissociation constant of multivalent RNA-binding proteins, here as an example for three binding sites on the RNA and the protein, based on the $K_\mathrm{d}$s of the individual domains. This system has 7 different RNA-protein complexes. According to the law of mass action we can write the total apparent $K_\mathrm{d}$ in terms of the concentrations of all individual states as

$$K_{\mathrm{d,\ tot}}(3)^{-1} = K_{\mathrm{a,\ tot}}(3) = \frac{[100]+[010]+[001]+[110]+[101]+[011]+[111]}{[000][\mathrm{RNA}]}\,. \quad (1)$$

This coupled reaction system has 12 individual equilibrium reactions (Figure 2B, main text), which can be described by their individual dissociation constants by applying the law of mass action, so that for example

$$K_{\mathrm{a}1} = \frac{[100]}{[000][\mathrm{RNA}]}\,, \qquad K_{\mathrm{a}2} = \frac{[010]}{[000][\mathrm{RNA}]}\,, \qquad K_{\mathrm{a}3} = \frac{[001]}{[000][\mathrm{RNA}]}\,, \quad (2)$$

$$K_{\mathrm{a},(100,110)} = \frac{[110]}{[100]} = K_{\mathrm{a}2} c_{d_1,L_1}\,, \qquad K_{\mathrm{a},(010,110)} = \frac{[110]}{[010]} = K_{\mathrm{a}1} c_{d_1,L_1}\,, \quad (3)$$

$$K_{\mathrm{a},(011,111)} = \frac{[111]}{[011]} = K_{\mathrm{a}1} c_{d_1,L_1}\,, \qquad K_{\mathrm{a},(110,111)} = \frac{[111]}{[110]} = K_{\mathrm{a}3} c_{d_2,L_2} \quad (4)$$

and similarly for all other reactions. The concentrations $c_{d_i,L_i}$ are expressed as described in the main text according to the Gaussian limit of the worm like chain model.

To express equation (1) in terms of known quantities we first plug in equations (2) to get

$$K_{\mathrm{a,\ tot}}(3) = K_{\mathrm{a1}} + K_{\mathrm{a2}} + K_{\mathrm{a3}} + \frac{[110] + [101] + [011] + [111]}{[000][\mathrm{RNA}]}\,. \tag{5}$$

By comparing equation (2) with equation (3) we get

$$\frac{[110]}{[010]} = \frac{[100]c_{d_1,L_1}}{[000][\mathrm{RNA}]}\,, \tag{6}$$

We can now expand this fraction with [110] and rearrange so that it becomes

$$\frac{[110]}{[000][\mathrm{RNA}]} = \frac{[110][110]}{[100][010]c_{d_1,L_1}}\,, \tag{7}$$

which reduces to

$$\frac{[110]}{[000][\mathrm{RNA}]} = K_{\mathrm{a1}}K_{\mathrm{a},(100,110)} = c_{d_1,L_1}K_{\mathrm{a1}}K_{\mathrm{a2}}\,. \tag{8}$$

With the same treatment we also get

$$\frac{[101]}{[000][\mathrm{RNA}]} = c_{|\vec{d_1}+\vec{d_2}|,L_1+L_2}K_{\mathrm{a1}}K_{\mathrm{a3}}\,, \quad \frac{[011]}{[000][\mathrm{RNA}]} = c_{d_2,L_2}K_{\mathrm{a2}}K_{\mathrm{a3}}\,. \tag{9}$$

Substituting equation (8) and equations (9) into (5) gives us

$$K_{\mathrm{a,\ tot}}(3) = K_{\mathrm{a1}}+K_{\mathrm{a2}}+K_{\mathrm{a3}}+c_{d_1,L_1}K_{\mathrm{a1}}K_{\mathrm{a2}}+c_{|\vec{d_1}+\vec{d_2}|,L_1+L_2}K_{\mathrm{a1}}K_{\mathrm{a3}}+c_{d_2,L_2}K_{\mathrm{a2}}K_{\mathrm{a3}}+\frac{[111]}{[000][\mathrm{RNA}]}\,. \tag{10}$$

To also express the last summand in terms of the individual dissociation constants, equation (4) can be compared against equation (2) giving

$$\frac{[111]}{[011]} = \frac{[100]c_{d_1,L_1}}{[000][\mathrm{RNA}]}\,. \tag{11}$$

Expanding the fraction with [111] and [110] yields

$$\frac{[111][111][110]}{[011]} = \frac{[100][110][111]c_{d_1,L_1}}{[000][\mathrm{RNA}]}\,. \tag{12}$$

Rearranging leads to

$$\frac{[111]}{[000][\mathrm{RNA}]} = \frac{[111]}{[011]c_{d_1,L_1}} \cdot \frac{[110]}{[100]} \cdot \frac{[111]}{[110]}\,, \tag{13}$$

which simplifies to

$$\frac{[111]}{[000][\mathrm{RNA}]} = c_{d_1,L_1}c_{d_2,L_2}K_{\mathrm{a1}}K_{\mathrm{a2}}K_{\mathrm{a3}}\,. \tag{14}$$

Plugging this into equation (10) gives us the total apparent $K_\mathrm{a}$ for cooperative binding with three binding sites as

$$K_{\mathrm{a,\ tot}}(3) = K_{\mathrm{a1}} + K_{\mathrm{a2}} + K_{\mathrm{a3}} + c_{d_1,L_1} K_{\mathrm{a1}} K_{\mathrm{a2}} + c_{|\vec{d_1}+\vec{d_2}|,L_1+L_2} K_{\mathrm{a1}} K_{\mathrm{a3}} + c_{d_2,L_2} K_{\mathrm{a2}} K_{\mathrm{a3}} + c_{d_1,L_1} c_{d_2,L_2} K_{\mathrm{a1}} K_{\mathrm{a2}} K_{\mathrm{a3}}\,. \tag{15}$$

#### 2 Procedure for calculating a dissociation constant for "fuzzy" binding and $N = 2$

In analogy to the above described treatment, we write the total $K_{\mathrm{a,\ tot}}$ for two domains, when we allow "fuzzy" binding (every domain can bind every other binding site), in terms of the concentrations of all binding configurations as

$$K_{\mathrm{a,tot}}(2) = \frac{[10][01][11][1_2 0][01_1][1_2 1_1]}{[00][\mathrm{RNA}]}\,, \tag{16}$$

where $[1_2 0]$ and $[01_1]$ denote the concentration of the first domain bound to the second RNA binding site, and the second domain bound to the first binding site, respectively. Again, we can write dissociation constants for individual binding steps. With simple substitutions we finally arrive at

$$K_{\mathrm{a,\ tot}}(2) = K_{\mathrm{a1}} + K_{\mathrm{a2}} + K_{\mathrm{a1_2}} + K_{\mathrm{a2_1}} + c_{d_1,L_1} K_{\mathrm{a1}} K_{\mathrm{a2}} + c_{d_1,L_1} K_{\mathrm{a1_2}} K_{\mathrm{a2_1}}\,, \tag{17}$$

where $K_{\mathrm{a1_2}}$ and $K_{\mathrm{a2_1}}$ denote the association constant for the first domain binding to the second RNA binding site, and the second domain binding to the first domain, respectively.

#### 3 Effect of flexible peptide linker on the effective concentration

If the protein binding domains are connected by flexible linkers, we need additional parameters to describe the local concentration, $c_{\mathrm{eff}}$ (Figure 1). It can be expressed by a combination of three distributions:

$$c_{R_1,R_2,L_{\mathrm{RNA}},L_{\mathrm{Protein}}} \approx \int\int \mathcal{N}\left(\vec{R_1} + \vec{r_p} + \vec{r_2} | 0, \sigma^2_{\mathrm{RNA}}\right) \mathcal{N}\left(\vec{r_p} | 0, \sigma^2_{\mathrm{Protein}}\right) \frac{\delta(\|\vec{r_2}\| - R_2)}{4\pi R_2^2} \mathrm{d}\vec{r_2}\mathrm{d}\vec{r_p} \tag{18}$$

with

$$\sigma^2_{\mathrm{RNA}} = \frac{2}{3} l_{p,\mathrm{RNA}} \cdot L_{\mathrm{RNA}} \quad \text{and} \quad \sigma^2_{\mathrm{Protein}} = \frac{2}{3} l_{p,\mathrm{Protein}} \cdot L_{\mathrm{Protein}}\,.$$

By integrating equation (18) over $\vec{r_p}$ and defining $\sigma^2 = \sigma^2_{\mathrm{RNA}} + \sigma^2_{\mathrm{Protein}}$, we can write it as

$$c_{R_1,R_2,L_{\mathrm{RNA}},L_{\mathrm{Protein}}} \approx \int \mathcal{N}\left(\vec{R_1} + \vec{r_2} | 0, \sigma^2\right) \frac{\delta(\|\vec{r_2}\| - R_2)}{4\pi R_2^2} \mathrm{d}\vec{r_2} \tag{19}$$

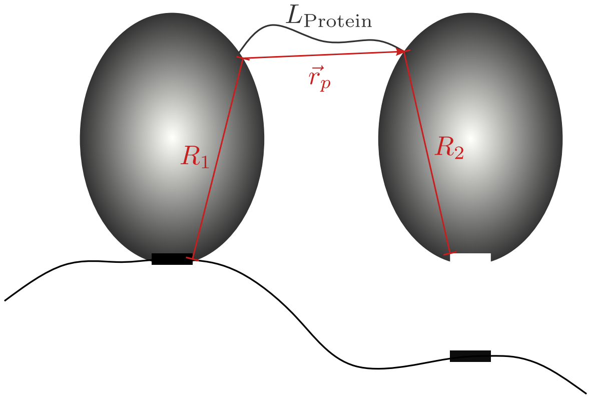


**Figure 1: Protein with flexible linker between domains and its RNA target.** To describe the effective concentration of the second RNA binding site at the second protein domain, we introduce $R_1$, $R_2$ and $L_{\text{Protein}}$ as new parameters.

To integrate over the spherical shell with radius $R_2$, we transform to spherical coordinates

$$c_{R_1,R_2,L_{\text{RNA}},L_{\text{Protein}}} \approx \frac{1}{4\pi R_2^2}\frac{1}{(2\pi\sigma^2)^{\frac{3}{2}}}\int_0^\pi \exp\left(-\frac{1}{2\sigma^2}\left(R_1^2+R_2^2-2R_1R_2\cos\theta\right)\right)2\pi R_2^2\sin\theta\mathrm{d}\theta \tag{20}$$

$$= \frac{1}{2(2\pi\sigma^2)^{\frac{3}{2}}}\exp\left(-\frac{1}{2\sigma^2}\left(R_1^2+R_2^2\right)\right)\int_0^\pi \exp\left(-\frac{R_1R_2\cos\theta}{\sigma^2}\right)\sin\theta\mathrm{d}\theta \tag{21}$$

$$= \frac{1}{2(2\pi\sigma^2)^{\frac{3}{2}}}\exp\left(-\frac{1}{2\sigma^2}\left(R_1^2+R_2^2\right)\right)\frac{\sigma^2}{R_1R_2}\left(\exp\left(\frac{R_1R_2}{\sigma^2}\right)-\exp\left(-\frac{R_1R_2}{\sigma^2}\right)\right) \tag{22}$$

and finally arrive at

$$c_{R_1,R_2,L_{\text{RNA}},L_{\text{Protein}}} \approx \frac{1}{(2\pi\sigma^2)^{\frac{3}{2}}}\frac{\exp\left(-\frac{1}{2\sigma^2}(R_1-R_2)^2\right)-\exp\left(\frac{1}{2\sigma^2}(R_1+R_2)^2\right)}{2R_1R_2/\sigma^2}\,. \tag{23}$$

### 4 Simulating cooperative binding with Gillespie stochastic simulation algorithm

The Gillespie algorithm allows us to define model parameters and reactions, and based on these obtain the time dependent concentrations of all species in the system after a simulation. To build the model we used the Python library Gillespy2. The parameters of the system are as described in the main text.

Before the simulation, all possible molecular entities in the system are defined, which in our case correspond to the different binding configurations. Furthermore, we define all possible reactions between these configurations with rate constants, where the reactions are binding and unbinding events. Because only the ratio between rates determines the end state, the off-rate constant of individual domains $k_{\text{off}}$ is set to $k_{\text{off}} = 1$ and according to $K_{\text{d}} = \frac{k_{\text{off}}}{k_{\text{on}}}$ the respective on-rate constant is $k_{\text{on}} = \frac{1}{K_{\text{d}}}$. We assume the rate constants of the unbinding events to be independent of the current binding state. The on-rate constants are calculated based on the current state as $k^*_{\text{on}} = k_{\text{on}} c_{\text{eff}}$ with $k_{\text{on}}$ the on-rate constant for the individual domain.

$c_{\text{eff}}$ is expressed according to the worm-like-chain model as a Gaussian distribution. When looking at a system with more than 2 binding sites, there are going to be reactions where an unbound binding site is between two bound binding sites as seen in the reaction $101 \rightleftharpoons 111$. The effective concentration in this case has to be expressed according to the laws of probability as the normalized product of two Gaussians. For the normalized product of two Gaussians we get

$$c_{\text{eff}} = \frac{1}{\left(2\pi\sigma_{12}^2\right)^{\frac{3}{2}}} \exp\left(-\frac{(d-\mu_{12})^2}{2\sigma_{12}^2}\right) \tag{24}$$

with

$$\sigma_{12}^2 = \frac{\sigma_1^2\sigma_2^2}{\sigma_1^2+\sigma_2^2} \quad \text{and} \quad \mu_{12} = \frac{\mu_1\sigma_1^2 + \mu_2\sigma_2^2}{\sigma_1^2+\sigma_2^2}\,.$$

In our case we have $d = d_1$, $\mu_1 = 0$ and $\mu_2 = d_1 + d_2$ with $d_1$ and $d_2$ the distances between binding domains on the protein.

The $K_{\text{d}}$ is then calculated based on the appropriate concentrations after reaching equilibrium in the simulation.

### 5 Model parameters for the comparison of experimental measurements to our theoretical estimates

| | $K_1$/M | $K_2$/M | RNA/nt | Protein/aa | $d$/m | $R_1$/m | $R_2$/m |
|---|---|---|---|---|---|---|---|
| ZBP1 | $2 \times 10^{-6}$ | $1.1 \times 10^{-6}$ | 18 | 0 | $3.85 \times 10^{-9}$ | 0 | 0 |
| hnRNP A1 | $20.4 \times 10^{-6}$ | $6.8 \times 10^{-6}$ | 4 | 0 | $1.99 \times 10^{-9}$ | 0 | 0 |
| PTB34 | $2.1 \times 10^{-6}$ | $2 \times 10^{-6}$ | 30 | 0 | $2.48 \times 10^{-9}$ | 0 | 0 |
| IMP3 — RRM12, KH12 | $9 \times 10^{-6}$ | $4 \times 10^{-6}$ | 6 | 39 | 0 | $5.38 \times 10^{-10}$ | $2.54 \times 10^{-9}$ |
